# Stimulation of the Dorsal Root Ganglion using an Injectrode^®^

**DOI:** 10.1101/2021.08.16.456553

**Authors:** Ashley N Dalrymple, Jordyn E Ting, Rohit Bose, James K Trevathan, Stephan Nieuwoudt, Scott F Lempka, Manfred Franke, Kip A Ludwig, Andrew J Shoffstall, Lee E Fisher, Douglas J Weber

## Abstract

**Objective:** The goal of this work was to compare afferent fiber recruitment by dorsal root ganglion (DRG) stimulation using an injectable polymer electrode (Injectrode^®^) and a more traditional cylindrical metal electrode.

**Approach:** We exposed the L6 and L7 DRG in four cats via a partial laminectomy or burr hole. We stimulated the DRG using an Injectrode or a stainless steel electrode using biphasic pulses at three different pulse widths (80, 150, 300 μs) and pulse amplitudes spanning the range used for clinical DRG stimulation. We recorded antidromic evoked compound action potentials (ECAPs) in the sciatic, tibial, and common peroneal nerves using nerve cuffs. We calculated the conduction velocity of the ECAPs and determined the charge-thresholds and recruitment rates for ECAPs from Aα, Aβ, and Aδ fibers. We also performed electrochemical impedance spectroscopy measurements for both electrode types.

**Main Results:** The ECAP thresholds for the Injectrode did not differ from the stainless steel electrode across all primary afferents (Aα, Aβ, Aδ) and pulse widths; charge-thresholds increased with wider pulse widths. Thresholds for generating ECAPs from Aβ fibers were 100.0 ± 32.3 nC using the stainless steel electrode, and 90.9 ± 42.9 nC using the Injectrode. The ECAP thresholds from the Injectrode were consistent over several hours of stimulation. The rate of recruitment was similar between the Injectrodes and stainless steel electrode and decreased with wider pulse widths.

**Significance:** The Injectrode can effectively excite primary afferents when used for DRG stimulation within the range of parameters used for clinical DRG stimulation. The Injectrode can be implanted through minimally invasive techniques while achieving similar neural activation to conventional electrodes, making it an excellent candidate for future DRG stimulation and neuroprosthetic applications.

## INTRODUCTION

Over the last several decades, neurostimulation, which is a neuromodulation therapy that involves the electrical stimulation of neural structures, has emerged as a promising treatment for various neurological conditions, ranging from Parkinson’s disease to depression and chronic pain. In the United States, 25 – 100 million people are affected by daily chronic pain (Institute of Medicine, 2011; Nahin, 2015), with approximately 10-20% suffering chronic pain from neuropathic origins (Bouhassira et al., 2008; van Hecke et al., 2014). The consequences of chronic pain go beyond physical pain; chronic pain also causes emotional distress, social isolation, and even career and financial strain (Hylands-White et al., 2017). The treatment of chronic pain often involves medications including analgesics, non-steroidal anti-inflammatory drugs, and opioids. Pharmacological treatment of chronic pain can be ineffective for a large number of patients and has greatly contributed to the opioid epidemic (Deyo et al., 2015; Marshall et al., 2019). Chronic pain treatment can also be achieved using invasive and permanent surgical procedures, including nerve blocks, dorsal root entry zone lesioning, and rhizotomies (Hylands-White et al., 2017). Neurostimulation may reduce the required medication dose and reduce the need for permanent surgical interventions (Deer et al., 2013).

A common neurostimulation therapy for chronic pain is epidural spinal cord stimulation (SCS), which utilizes electrodes that are implanted in the dorsal epidural space to electrically stimulate the dorsal columns and other structures within the spinal cord (Caylor et al., 2019). Presently, an estimated 50,000 SCS devices are implanted every year (Sdrulla et al., 2018). In recent years, the dorsal root ganglion (DRG) has emerged as a new target for neurostimulation (Deer et al., 2013; Liem et al., 2013). A device to stimulate the DRG was approved for clinical use in the United States in 2016 (Deer et al., 2019).

Several hypotheses have been proposed to explain the mechanisms of both SCS and DRG stimulation. One common hypothesis is the gate control theory of pain (Melzack and Wall, 1965), in which neurostimulation activates Aβ primary afferent fibers, which then activate inhibitory interneurons in the substantia gelatinosa of the dorsal horn, reducing the firing of wide dynamic range neurons that also respond to Aδ- and C-fiber activation (Caylor et al., 2019; Graham et al., 2019). Therefore, measuring the stimulation threshold to activate Aβ fibers is relevant to evaluating neurostimulation devices for treating chronic pain.

The DRG is an advantageous target for neurostimulation because it offers precise coverage of a dermatome corresponding to the painful region (Esposito et al., 2019; Liem, 2015). Multiple clinical trials have shown that DRG stimulation is effective in treating chronic pain stemming from various sources, including back pain, distal limb pain, phantom limb pain, complex regional pain syndrome, causalgia, pelvic pain, groin pain, and mononeuropathies (Eldabe et al., 2015; Harrison et al., 2018; Hunter and Yang, 2019; Van Buyten et al., 2015; Zuidema et al., 2014). A recent clinical trial comparing the effectiveness of SCS and DRG stimulation for the treatment of complex regional pain syndrome and causalgia showed that DRG stimulation produced a greater than 50% reduction in pain in 81.2% of participants, compared to only 55.7% of participants who received SCS (Deer et al., 2017).

A factor limiting adoption of DRG stimulation is the surgical approach (Deer et al., 2013). DRG lead delivery methods also differ by spinal level. The surgical technique for placing DRG leads targeting the thoracic and lumbar levels involves implanting the leads into the epidural space and steering them laterally through the neuroforamen and placed near the DRG (Caylor et al., 2019; Deer et al., 2019; Liem, 2015). When targeting the sacral DRG, however, the leads are placed using a transforaminal approach (Deer et al., 2019). Using the current lead systems, the transforaminal approach is more likely to result in lead migration than the epidural approach (Deer et al., 2019). Furthermore, implanting DRG stimulation leads can be uncomfortable for patients and many permanent implants are often done under general anaesthesia (Deer et al., 2019; Morgalla et al., 2017). There has also been much debate surrounding the complication rates and adverse events related to DRG stimulation. While a recent report suggested DRG stimulation has an excellent safety record (Deer et al., 2020), several studies have reported adverse events, such as pain near the site of the implantable pulse generator and lead fracture (Deer et al., 2017; Horan et al., 2020; Huygen et al., 2020). Some experts believe that the complications are due to the increased time and learning curve required for performing the DRG stimulation lead implant procedure, as well as the recency of its clinical implementation (Lubenow and Nijhuis, 2020). An electrode that conforms to tissue and can still be placed in close proximity to the DRG using a minimally invasive procedure would reduce such barriers to adoption of a clinically efficacious treatment.

To this end, we have developed the Injectrode^®^, which is a flowable two-part conductive pre-polymer that can be injected through a needle and syringe into the space adjacent to the neuroanatomical target, and then quickly polymerizes *in vivo*, creating a highly conformal functioning electrode. The version of the Injectrode evaluated here follows previous work, which showed that a conductive polymer Injectrode formulation remains soft and flexible after curing, with a Young’s modulus that more closely matches that of the tissue compared to traditional metal electrodes (Trevathan et al., 2019). Soft electrodes have been shown in other studies to reduce the inflammatory response and improve electrode acceptance (Sohal et al., 2016). Trevathan and colleagues also included initial proof of concept data demonstrating that the Injectrode can be used for stimulation of the brachial plexus of rats to produce tetanic muscle contractions, and stimulation of the vagus nerve in swine to induce a reduction in heart rate (Trevathan et al., 2019). However, a comprehensive characterization of the Injectrode in terms of recruitment profiles of nerve fiber types associated with therapeutic effect was not performed. Ongoing work is aimed at developing and testing the Injectrode to ensure that it remains soft and flexible, can effectively stimulate neural tissue, can be delivered using a simple and minimally invasive procedure (such as a transforaminal approach), and to utilize a collector for transcutaneous stimulation to reduce the need for an implantable pulse generator (IPG) and likelihood of lead migration. A simplified system using the Injectrode may reduce uptake barriers, leading to the DRG stimulation being used at an earlier stage of treatment instead of a last resort.

The goal of the current study was to investigate if the Injectrode is a viable electrode for DRG stimulation. We evaluated the ability of the Injectrode to recruit primary afferents during DRG stimulation in cats, as these afferents, particularly Aβ fibers, are thought to drive the therapeutic effects of DRG stimulation for the treatment of pain. We compared charge thresholds for recruiting primary afferent fibers using the Injectrode and a cylindrical stainless steel electrode placed on the surface of the DRG. A stainless steel electrode was chosen as a surrogate for a clinical DRG lead because it is a well characterized material, has similar shape and stiffness to clinical leads, is relatively inexpensive to purchase, and readily accessible for proof-of-concept testing.

We demonstrated that the Injectrode can be used to stimulate a range of afferent fibers, including Aα, Aβ, and Aδ fibers, in the DRG with charge thresholds similar to or lower than a cylindrical stainless steel electrode. Additionally, both electrodes effectively recruited Aβ fibers, the putative target of DRG stimulation, across a range of stimulation parameters, including current amplitude, pulse width, and frequency, within the range typically used for clinical DRG stimulation. Although further work is needed for validation, this study illustrates that the Injectrode may offer an alternative to standard clinical DRG stimulation electrodes. As an injectable and conformal material, the Injectrode may offer important advantages, including ease of delivery and greater stability of contact with the DRG.

## METHODS

### Experimental Model

We performed acute experiments in four adult male cats (4.38-7.55 kg). All procedures were approved by the University of Pittsburgh Institutional Animal Care and Use Committee. We induced anaesthesia using ketamine (cat A; intramuscular, 10 mg/kg) and acepromazine (cat A; intramuscular, 0.1 mg/kg), or dexdomitor (cats B-D; intramuscular, 0.04 mg/kg). We then intubated the cat and administered isoflurane for the duration of the experiment (inhalation, 2-2.5%). We used antisedan (intramuscular; 1:1 dexdomitor dose) to reverse the effects of dexdomitor following the administration of isoflurane. We gave atropine intramuscularly (all cats; 0.05 mg/kg) to reduce saliva output. We continuously monitored vital signs including heart rate, SpO_2_, core temperature, respiratory rate, and ETCO_2_. We also continually administered IV fluids (saline, 0.9% sodium chloride) throughout the experiment.

### Manufacturing of Dorsal Root Ganglion Electrodes

Injectrodes were manufactured by Neuronoff, Inc. (Cleveland, OH, USA) using a similar polymer-conductor variant of the Injectrode as described previously by Trevathan et al., 2019. Briefly, we mixed two parts of Pt-curing silicone elastomers (World Precision Instruments, FL, USA) with metallic silver particles (Sigma-Aldrich, MO, USA) and loaded the mixture into a syringe that was calibrated to hold 10 μL of Injectrode material. Silver was chosen for this proof-of-concept study of a particle-polymer-based electrode (the Injectrode) because it was readily available and cost-efficient. An insulated silver wire (AS-766-36; Cooner Wire Company, Chatsworth, CA, USA) with de-insulated ends was placed inside the syringe such that it became embedded in the Injectrode material upon curing on one end, while the other end was connected to the stimulator for DRG stimulation (see below). Based on measurements taken from explanted Injectrodes, the lengths and diameters of the Injectrodes ranged from 4.0 – 6.0 (4.4 ± 0.7) mm and 1.3 – 3.1 (1.8 ± 0.6) mm, respectively. These parameters correspond to an average surface area of 30.8 ± 12.5 mm^2^. We manufactured stainless steel DRG electrodes in-house (University of Pittsburgh) using 1 mm diameter hypodermic tubing (McMaster-Carr, Elmhurst, IL, USA) cut to a length of 4.5 mm and crimped to a Teflon-insulated lead wire (AS632; Cooner Wire Company, Chatsworth, CA, USA). The surface area of the stainless steel electrode was 15.7 mm^2^, which is 1.96 times smaller than the average surface area of the Injectrode.

### Implantation of Electrodes

We placed the DRG electrodes on the L6 and L7 DRG after exposing the DRG via partial laminectomy or burr hole in the vertebral laminae. In cat A, we performed a partial laminectomy to expose both target DRGs, and in cat B, only the L7 DRG was exposed via a partial laminectomy. For the laminectomy exposures, we placed the electrodes on the dorsal surface of the crown of the DRG for stimulation. We filled the open cavity with 5 mL of saline to keep the tissue from drying. For the L6 DRG in cat B, as well as both the L6 and L7 DRG in cats C and D, we drilled a burr hole through the laminae overlying the DRG (Figure 1). For the burr hole exposure, we placed the stainless steel electrode vertically in the hole, while the Injectrode was injected into the hole by placing the tip of the blunt needle into the hole (Supplementary Video 1). The placement of the burr hole was guided by the exposing and matching the locations of the DRG on the contralateral side, which were exposed via a partial laminectomy. Following testing with the electrodes delivered through the burr hole, we then removed the lamina to confirm the location of the Injectrodes. In all experiments, we placed a stimulation ground electrode (AS636, Cooner Wire Company, Chatsworth, CA, USA) with ~5 cm exposure in between the skin and lumbodorsal fascia ipsilateral to the DRG electrodes for monopolar stimulation.

**Figure 1.**
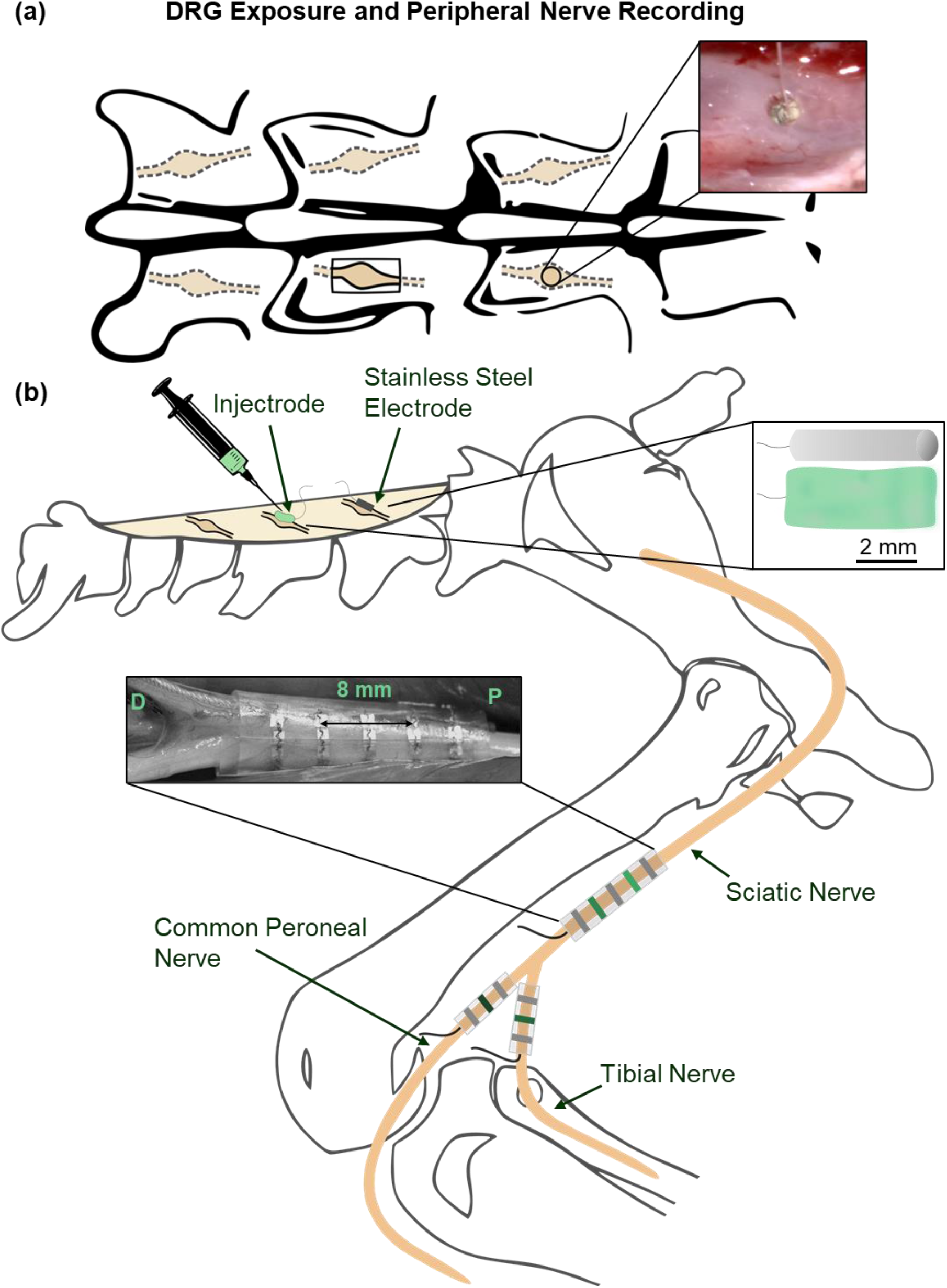
Experimental setup. (a) Exposure of dorsal root ganglion (DRG) by either a partial laminectomy or a burr hole. Inset: Injectrode delivered through a burr hole over the DRG. (b) An Injectrode or a stainless steel electrode were placed on top of a DRG for delivering stimulation. Antidromic evoked compound action potentials were recorded using spiral nerve cuffs placed on the sciatic, tibial, and common peroneal nerves. Inset: five-contact nerve cuff placed on sciatic nerve. Cuff contacts were 4 mm apart. Gray nerve cuff contacts (1, 3, 5 for sciatic or 1, 3 for tibial and common peroneal) were shorted together and used as a reference. Green nerve cuff contacts (2, 4 for sciatic or 2 for tibial and common peroneal) were used for recording. P= proximal; D = distal. Inset: Scaled illustration of the stainless steel electrode and Injectrode.

We implanted three spiral nerve cuff electrodes made of silicone with platinum contacts in the hind limb ipsilateral to the stimulated DRG (Figure 1; Ardiem Medical, Indiana, PA, USA). A five-contact spiral nerve cuff electrode (3 mm diameter, contacts 4 mm apart, center-to-center) was implanted around the sciatic nerve. The proximal, center, and distal contacts were shorted together and used as a reference electrode. The second and fourth contacts were used for recording and, with the references, created two tripolar configurations. Spiral nerve cuff electrodes with three contacts (2 mm diameter, contacts 4 mm apart, center-to-center) were implanted around the tibial and common peroneal nerves. The proximal and distal contacts were shorted together to create a tripolar configuration with the center electrode used for recording. The nerve cuffs recorded the compound action potentials evoked by stimulation of the DRG.

### Data Acquisition and Stimulation Protocol

We recorded electroneurogram (ENG) signals using a Grapevine Neural Interface Processor (Ripple, Salt Lake City, UT, USA) and a single-ended headstage (Surf S2), which has an input range of ±8 mV and 16-bit resolution with 0.25 μV/bit. We filtered the ENG using a 0.1 Hz high pass filter and a 7.5 kHz low pass filter, followed by digitization at 30 kS/s.

We delivered monopolar stimulation using a high current ECOG + Stim front end (Ripple, Salt Lake City, UT, USA; Cats A and B) or an A-M Systems 4100 Stimulator (A-M Systems, Sequim, WA, USA; Cats C and D). Stimulation was delivered at 58 Hz using biphasic, symmetric, cathodic-leading pulses. Stimulation pulse widths were either 80, 150, or 300 μs per phase. In cat A, only 80 and 150 μs pulse widths were tested. At each DRG location and for both stainless steel electrodes and Injectrodes, we constructed recruitment curves by varying the stimulation amplitude from below sensory threshold up to motor threshold in steps of 20 - 250 μA (for coarse and fine steps). We determined motor threshold by delivering a train of five stimulation pulses and observing twitches in the hindlimb. The order of stimulation amplitudes in the recruitment curve was randomized, with 600 pulses repeated at each amplitude, and 5 – 10 s pauses between changes in amplitude. We repeated this process for each pulse width.

### ENG Analysis

We conducted all analyses using custom written programs in MATLAB (MathWorks, Natick, MA, USA). During experiments, we visualized responses online to select the stimulation amplitude range for the recruitment curves. We interpolated over a 1-ms window to blank stimulation artefacts. We filtered the ENG using a second-order high-pass filter with a cutoff frequency of 300 Hz, and we stimulation-triggered averaged the responses to 600 stimulus pulses. We determined the presence of an ECAP at a particular stimulation amplitude by comparing the root-mean-square (RMS) amplitude of the ENG to a threshold value, as shown in prior work (Ayers et al., 2016; Nanivadekar et al., 2019). We used a bootstrapping method to verify consistency of the evoked responses. For each stimulation amplitude, we generated a stimulus-triggered average of the ENG on each nerve cuff electrode using a random sample of 80% of the 600 total stimulus repetitions. This step was repeated 200 times. The averaged waveforms were then rectified and smoothed by calculating the RMS of each averaged ENG signal using a 100-μs sliding window with a 33-μs overlap with the previous window. From each averaged RMS ENG signal, we defined the ECAP threshold as one standard deviation above the upper limit of the 99% confidence interval of the baseline RMS amplitude during a 1-ms period before each stimulation pulse. An ECAP was detected if 95% of the subsets of random samples were supra-threshold during the same 100-μs window of the RMS ENG signal. Since we used a total of four nerve cuff electrodes (two on the sciatic plus one each on the tibial and common peroneal nerves), we defined threshold to be the lowest stimulation amplitude that evoked a response in any of the four nerve cuff electrodes. We confirmed the accuracy of the automated approach by visually identifying stimulation thresholds.

We calculated the conduction velocity of the ECAPs using the ENG from the sciatic nerve cuffs by following a procedure described previously by our group (Fisher et al., 2014). Briefly, we measured the time difference between the peaks of the first, short-latency ECAP recorded from the proximal and distal tripoles in the five-contact cuff (contacts 2 and 4; Figure 2). Contacts 2 and 4 are 8 mm apart, center-to-center. We divided that distance by the time difference (dt) between peaks of the ECAPs to obtain the conduction velocity for the lowest latency ECAP. If multiple ECAPs were present, we calculated the conduction velocity of the longer latency ECAP(s) by first determining the distance between the DRG electrode and the proximal tripole using the conduction velocity of the first ECAP and the latency between the onset of the stimulus and the peak of that ECAP. Then, we divided the distance between the stimulating and recording electrodes by the latency of the later ECAP(s) to obtain their conduction velocity.

**Figure 2.**
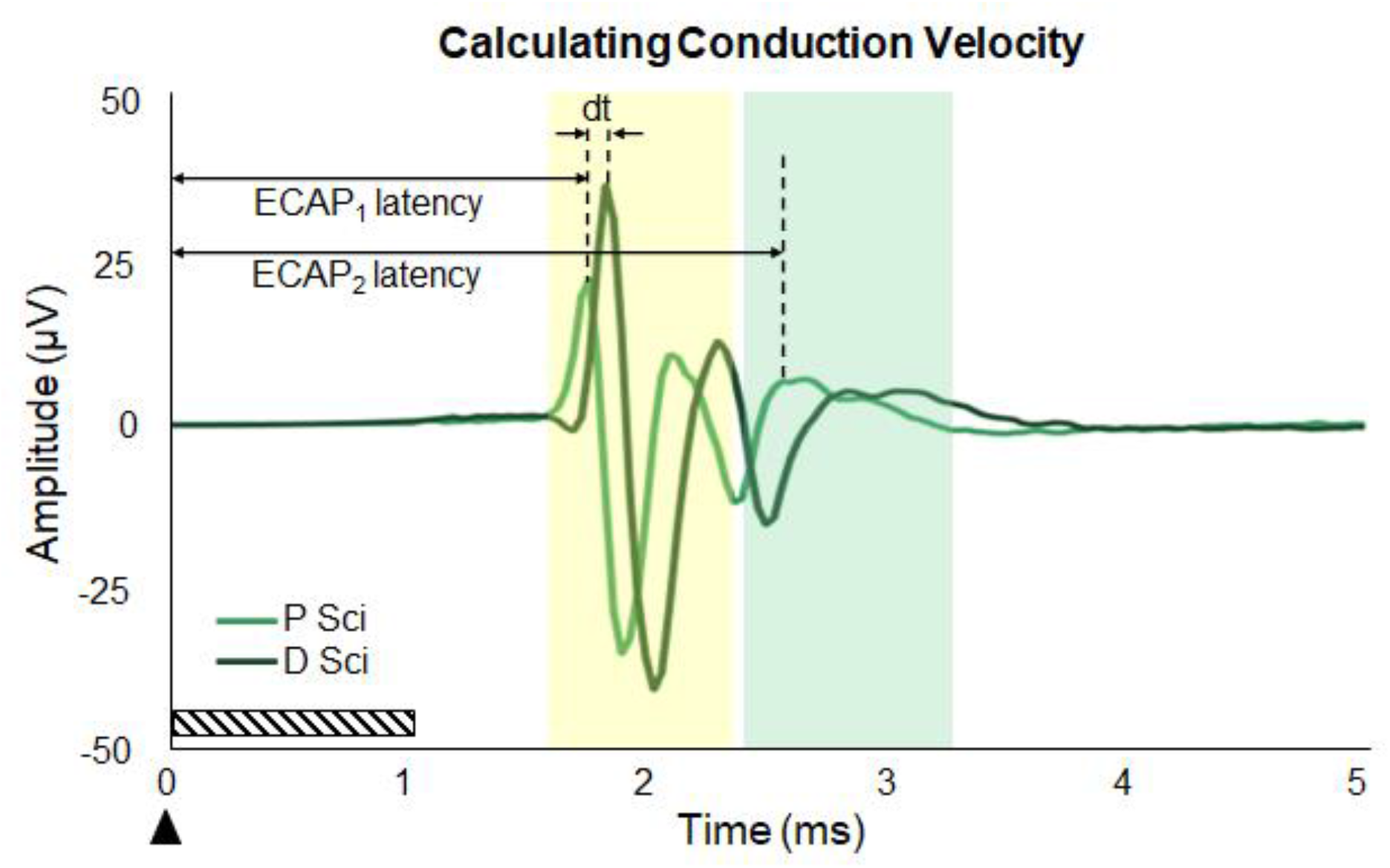
Calculating conduction velocity. Evoked compound action potentials (ECAPs) from the proximal (P Sci; contact 2) and distal (D Sci; contact 4) recording sites on the sciatic nerve cuff were aligned to the stimulation onset (arrow). The stimulation artefact was blanked in the region shown by the hashed bar. The time difference between the peaks of each ECAP was measured (dt). The two recording electrodes in the nerve cuff were 8 mm apart. That distance, along with dt, were used to calculate the conduction velocity of the first ECAP. Then, that conduction velocity and the latency of the first ECAP were used to calculate the distance between the stimulation and recording sites. Finally, the distance between the stimulation and recording sites, and the latency of the second ECAP, were used to determine the conduction velocity of the second ECAP.

Using the calculated distance between the DRG electrode and cuff electrode, and the known conduction velocity ranges for primary afferent axons, Aα (80-120 m/s), Aβ (35-80 m/s), or Aδ fibers (5-35 m/s) (Bear et al., 2007), we also defined latency windows from the onset of stimulation for each fiber type. The latency windows corresponding to the conduction velocities of the primary afferents were approximately 1.3 - 2.2 ms (Aα), 2.2 - 6.5 ms (Aβ), and 6.5 - 45 ms (Aδ), varying slightly according to the distance between the stimulation and recording electrodes. Using those latency windows, we found the ECAP threshold for the different fiber types (Aα, Aβ, and Aδ) and constructed recruitment curves for Aα and Aβ thresholds and each pulse width and depicted the peak-to-peak amplitude of the ECAP as a function of stimulation charge (current amplitude × pulse width).

We determined the recruitment rate for Aα and Aβ fibers by fitting a line to the linear region of the recruitment curves using least squares regression (MATLAB’s polyfit function, first order). Recruitment curves were not constructed for Aδ fibers because there were not enough data points for each condition; ECAPs from Aδ fibers were only detected when stimulation amplitude was very high (close to motor threshold). The recruitment rate is the slope of the best-fit line. To compare the recruitment rates across all trials, we normalized all slope values to the slope of the recruitment curve obtained for the stainless steel electrode and a pulse width of 300 μs, which is the pulse width used typically in clinical applications of DRG stimulation (Deer et al., 2017; Graham et al., 2019; Kent et al., 2018; Liem et al., 2013).

### Electrochemical Impedance Spectroscopy

We conducted *in vivo* electrochemical impedance spectroscopy (EIS) measurements of the stainless steel electrode and Injectrode on the L7 DRG contralateral to those used for stimulation in cats C-D at the beginning of each experiment, as well as for the Injectrodes that were left in the insertion holes in cat D following all stimulation trials. We used a three-electrode setup, where the working electrode was the DRG stimulation electrode (either stainless steel or Injectrode), the counter electrode was a Platinum wire (CHI115, CH Instruments, Inc., Austin, TX, USA; 32 mm long, 0.5 mm diameter) placed distally in between the skin and muscle of the leg ipsilateral to the active electrode, and the reference electrode was a Ag|AgCl wire (CHI111, CH Instruments, Inc., Austin, TX, USA; 40 mm long, 0.5 mm diameter) placed in between the skin and muscle of the back near the active electrode. We used a potentiostat (CompactStat, Ivium Technologies, Eindhoven, The Netherlands) to perform EIS measurements. EIS frequency values ranged from 1 to 100,000 Hz at eight points per decade and a peak-to-peak voltage of 25 mV.

### Statistical Methods

We used the Shapiro–Wilk test to test for normality and Levene’s test to assess the homogeneity of variance. We used a student’s t-test to compare threshold and recruitment rate means between the L6 and L7 DRG. We performed a two-way analysis of variance (ANOVA) to test for effects of material (Injectrode or stainless steel) and fiber type (Aα, Aβ, or Aδ) on ECAP thresholds. We also performed two-way ANOVAs to test for significant effects of material and pulse width (80, 150, or 300 μs) on ECAP thresholds and recruitment rate. We used the Bonferroni correction for multiple comparisons. ANOVA and the post-hoc tests were performed using SPSS Statistics (version 26; IBM, Armonk, NY, USA). We used a paired t-test to compare ECAP thresholds over time using Excel (version 2108; Microsoft Corporation, Redmond, WA, USA). We performed equivalence testing using the two one-sided t-test (TOST) method (Lakens, 2017) in MATLAB using the TOST function (Rastogi, 2017) to compare the thresholds of the Injectrode versus the stainless steel electrode across fiber types and over time. We considered a p-value ≤ 0.05 to indicate significance. Results are presented as mean ± standard deviation.

## RESULTS

We stimulated the L6 and L7 DRG in four cats using either a stainless steel electrode or an Injectrode. We recorded ECAPs in sensory nerve fibers measured using nerve cuff electrodes placed on the sciatic, tibial, and common peroneal nerves. We placed stimulation electrodes on the DRG after exposure via partial laminectomy or a burr hole. For all performance metrics, there were no significant differences in recruitment properties between the partial laminectomy exposure and the burr hole exposure. Therefore, we grouped these data together for all analyses.

Following data collection, we explanted several of the Injectrodes and measured their widths and lengths. Injectrodes delivered on the DRG exposed with a partial laminectomy were cylindrical in shape, having been formed by the needle used for delivery. Injectrodes that were injected into the burr-hole were still somewhat cylindrically shaped, but flatter as they filled the cavity between the DRG and the bone.

### DRG stimulation excites multiple types of primary afferent neurons

DRG stimulation is thought to induce analgesic effects through the activation of afferent fibers. Here, DRG stimulation generated ECAPs at multiple latencies in each of the nerve cuff recordings (Figure 3a), indicating recruitment of multiple fiber types. By measuring the propagation delay between ECAPs recorded at the two tripolar recording sites in the sciatic nerve cuff, we calculated the conduction velocities of the ECAPs and used them to classify the response as from Aα (80-120 m/s), Aβ (35-80 m/s), or Aδ (5-35 m/s) fibers (Bear et al., 2007). Evoked responses included conduction velocities ranging from 13 m/s to 120 m/s, which includes three primary afferent fiber types. Our method did not detect any C fiber responses.

**Figure 3.**
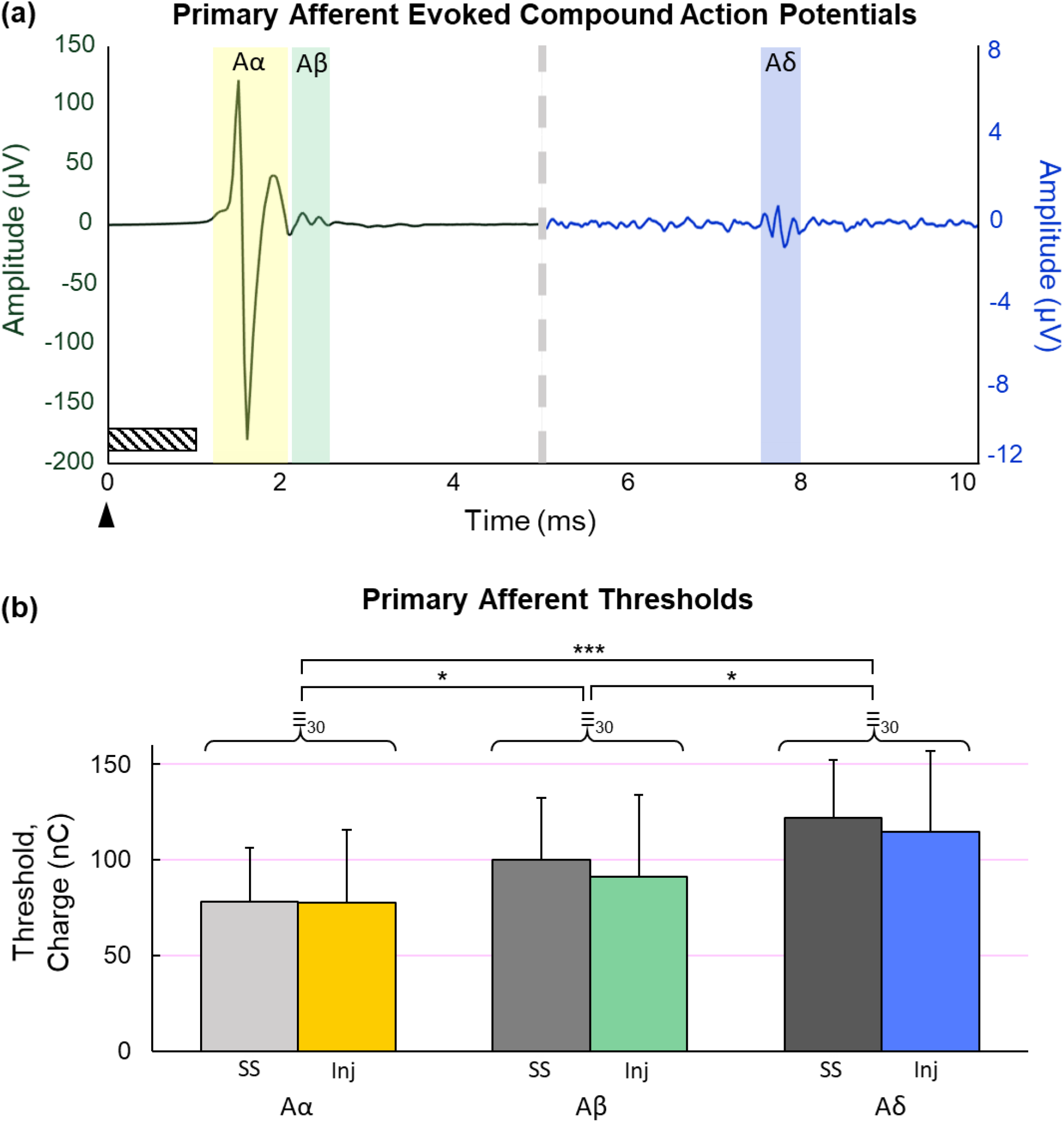
DRG stimulation evoked responses in primary afferents. (a) Examples of evoked compound action potentials (ECAPs) for Aα, Aβ, and Aδ fibers (highlighted). The stimulation onset is indicated by an arrow and the artefact was blanked (indicated by the hashed region). The ECAPs from the Aα and Aβ fibers are scaled according to the left y-axis; the ECAP from the Aδ fiber is scaled according to the right y-axis; the change in scales is indicated by the vertical dashed gray line (~5 seconds). (b) Stimulation thresholds required to elicit ECAPs from Aα, Aβ, and Aδ fibers using either the stainless steel (SS) electrode or the Injectrode (Inj). *p ≤ 0.05; ***p ≤ 0.001. ≡_30_ indicates that the thresholds of the Injectrode were equivalent to the thresholds of the stainless steel electrode within a ±30% window.

Aα fibers had the lowest threshold (77.8 ± 33.3 nC; Figure 3b). The threshold for recruiting Aβ fibers was 23% higher than the Aα threshold (95.4 ± 37.9 nC), which was significant (p = 0.034). The threshold for recruiting Aδ fibers was 118.0 ± 36.8 nC, which was 52% higher than the Aα threshold (p < 0.001) and 24% higher than the Aβ threshold (p = 0.033). We found no significant differences between the ECAP thresholds for the Injectrode versus the stainless steel electrode across all primary afferent fiber types (p = 0.412). The thresholds from the Injectrode were equivalent to the thresholds from the stainless steel electrode within a ±30% window for Aα (p = 0.007), Aβ (p = 0.024), and Aδ (p = 0.031) fibers.

### The Injectrode^®^ and stainless steel electrodes have similar ECAP thresholds that vary by pulse width

ECAP thresholds were significantly higher when stimulation was delivered to L7 compared to L6 for Aα and Aβ fibers (p < 0.001). There was no difference between the L6 and L7 thresholds for Aδ fibers (p = 0.63). Because there were differences in the Aα and Aβ fiber thresholds, we kept these analyses separate for each DRG level.

Differences in ECAP thresholds due to pulse width were not consistent across DRG levels. For Aα fibers, there were no significant differences between ECAP thresholds when stimulation was applied with a pulse width of 80 μs (49.0 ± 19.8 nC), 150 μs (57.8 ± 21.5 nC), or 300 μs (65.6 ± 29.3 nC) on the L6 DRG (Figure 4a; p = 0.34). However, on the L7 DRG, thresholds corresponding to stimulation with a 300-μs (124.9 ± 24.0 nC) pulse width were significantly higher than those corresponding to stimulation with an 80-μs (76.7 ± 22.4 nC; p < 0.001) pulse width, but not a 150-μs pulse width (97.7 ± 20.6 nC; p = 0.063). In general, the ECAP threshold increased with a longer pulse width. Notably, the ECAP thresholds for Aα fibers for the Injectrode were not significantly different from thresholds for stainless steel electrodes at L6 (Injectrode = 49.6 ± 22.4 nC; stainless steel = 63.7 ± 23.4 nC; p = 0.12) or L7 (Injectrode = 100.4 ± 32.2 nC; stainless steel = 93.3 ± 25.3 nC; p = 0.52). There were also no interaction effects between electrode type and pulse width (L6: p = 0.98; L7: p = 0.99).

**Figure 4.**
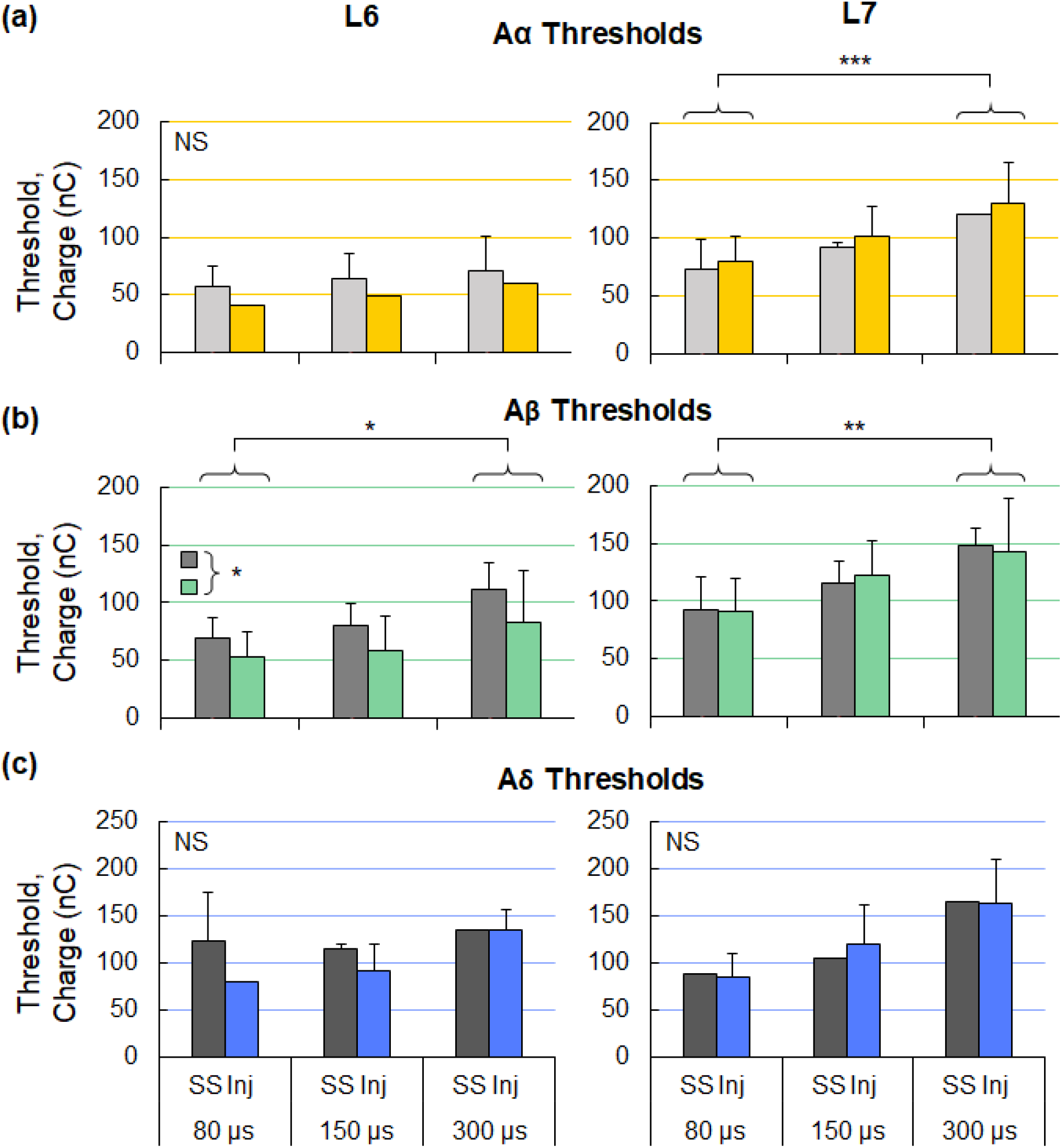
Stimulation thresholds to elicit evoked compound action potentials (ECAPs) for the L6 and L7 DRG for (a) Aα, (b) Aβ, and (c) Aδ fibers, across pulse widths. *p ≤ 0.05; **p ≤ 0.01; ***p ≤ 0.001. NS = not significant.

The ECAP threshold for Aβ fibers was significantly higher for the longest versus shortest pulses (300 μs vs 80 μs, Figure 4b), and the threshold difference was slightly greater at L7 compared to L6. At the L6 DRG, stimulation with a pulse width of 80 μs (60.9 ± 21.0 nC) had a significantly lower threshold than stimulation at 300 μs (97.1 ± 35.7 nC; p = 0.028). Similarly, stimulation on the L7 DRG with a pulse width of 80 μs (91.8 ± 26.2 nC) had a significantly lower threshold than stimulation at 300 μs (145.7 ± 28.8 nC; p = 0.002). Similar to the Aα fibers, the ECAP threshold increased with a longer pulse width. The thresholds for evoking ECAPS in Aβ fibers using the Injectrode were significantly lower than with the stainless steel electrodes at L6 (Inj = 64.0 ± 32.6 nC; SS = 84.8 ± 25.7 nC; p = 0.049) but not L7 (Inj = 115.9 ± 36.0 nC; SS = 116.4 ± 31.4 nC; p = 0.95). There were no interaction effects between the electrode type and pulse width (L6: p = 0.91; L7: p = 0.91).

Finally, for Aδ fibers, the ECAP thresholds did not differ across pulse widths for the L6 DRG (p = 0.27) or the L7 DRG (Figure 4c; p = 0.14). Furthermore, ECAP thresholds between the Injectrode and stainless steel electrodes were not significantly different at the L6 (Inj = 103.8 ± 31.9 nC; SS = 122.9 ± 29.1 nC; p = 0.25) or L7 (Inj = 123.1 ± 48.8 nC; SS = 119.3 ± 40.5 nC; p = 0.90) DRG. There were also no interaction effects between the electrode type and pulse width (L6: p = 0.67; L7: p = 0.96). However, there is a general but insignificant trend of an increasing threshold for longer pulse widths.

Taken together, ECAP thresholds for each fiber type (Aα, Aβ, Aδ) for DRG stimulation delivered through an Injectrode were either not different or significantly lower than the ECAP thresholds from stimulation with a stainless steel electrode. Generally, the ECAP thresholds increased as pulse width increased, but this trend was not always significant.

### Injectrodes have similar ECAP thresholds over time

We compared the stability of ECAP thresholds for evoking responses in Aα and Aβ fibers using DRG stimulation between the stainless steel electrode and the Injectrode. We measured thresholds early in the experiment and repeated those measurements 2 to 10.5 hours later (Figure 5). Early thresholds for evoking a response in Aα fibers was not significantly different from the later thresholds for the stainless steel electrode (early = 106.7 ± 25.1 nC; late = 102.8 ± 29.7 nC; p = 0.45) or the Injectrode (early = 89.7 ± 38.9 nC; late = 85.6 ± 34.5 nC; p = 0.27). There were also no significant differences over time for the Aβ thresholds for the stainless steel electrode (early = 120.7 ± 31.6 nC; late = 118.2 ± 37.6 nC; p = 0.64) or the Injectrode (early = 98.1 ± 34.8 nC; late = 102.2 ± 44.1 nC; p = 0.48). The thresholds measured at early and late timepoints were equivalent within a ±35% window for both Aα (p = 0.031) and Aβ (p = 0.038) fibers using the stainless steel electrode. The thresholds measured at early and late timepoints using the Injectrode were equivalent within a ±40% window for both Aα (p = 0.035) and Aβ (p = 0.032) fibers. Therefore, the stainless steel electrode, and more importantly, the Injectrode, had stable ECAP thresholds over many hours and thousands of stimulation pulses.

**Figure 5.**
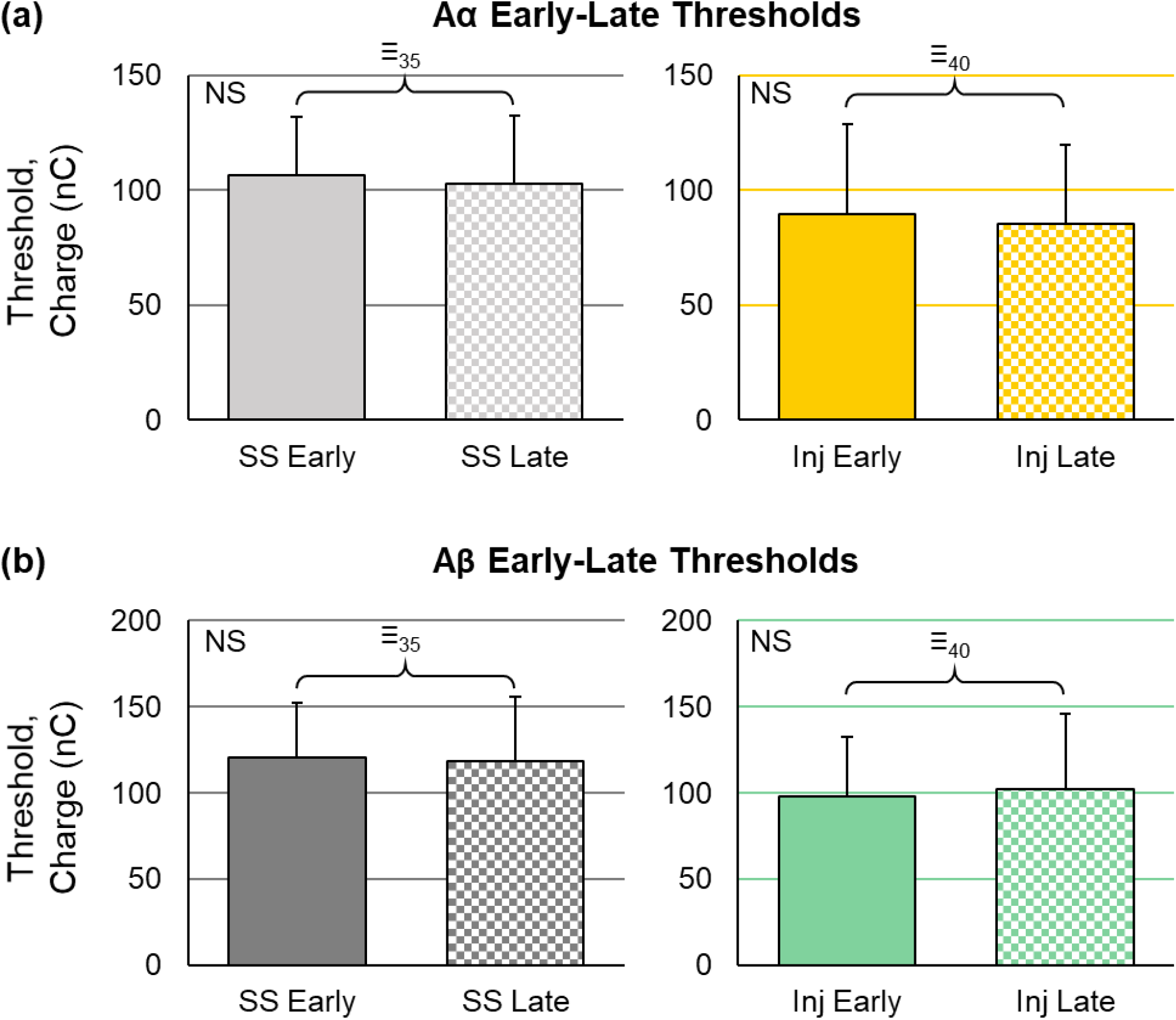
Stimulation thresholds to elicit evoked compound action potentials using the stainless steel (SS) electrode or the Injectrode (Inj) over time (2 to 10.5 hours) for (a) Aα and (b) Aβ fibers. NS = not significant. ≡35 and ≡40 indicate that the later thresholds were equivalent to the early thresholds within a ±35% and ±40% window, respectfully.

### Aα and Aβ fibers are more likely to be excited within the usable range than Aδ fibers

Clinical DRG stimulation generally operates within a current amplitude range above sensory threshold, where paresthesias are elicited, but below a level where any pain or movement is produced. We defined the usable range of DRG stimulation as the stimulation amplitudes between the ECAP threshold and just below the motor threshold. The ECAP threshold was defined as the lowest stimulation amplitude that evoked an ECAP, and the motor threshold was the stimulation amplitude that evoked a visible twitch in the lower limb. Data were combined for both electrode types to evaluate the mechanistic activation of afferent fibers using DRG stimulation. At the top of the usable range, or the maximum stimulation amplitude below motor threshold, ECAPs from Aα fibers were always present (100% of the trials), ECAPs from Aβ fibers were present in 85.7% of the trials, and ECAPs from Aδ fibers were present in 37.5% of the trials (Figure 6a). However, at the half-maximum point, which we defined as the stimulation amplitude mid-way between the ECAP threshold and top of the usable range, there were far fewer ECAPs from Aδ fibers detected (5.4% of the trials). ECAPs from Aα fibers (100% of trials) and Aβ fibers (75.0% of trials) remained prevalent.

**Figure 6.**
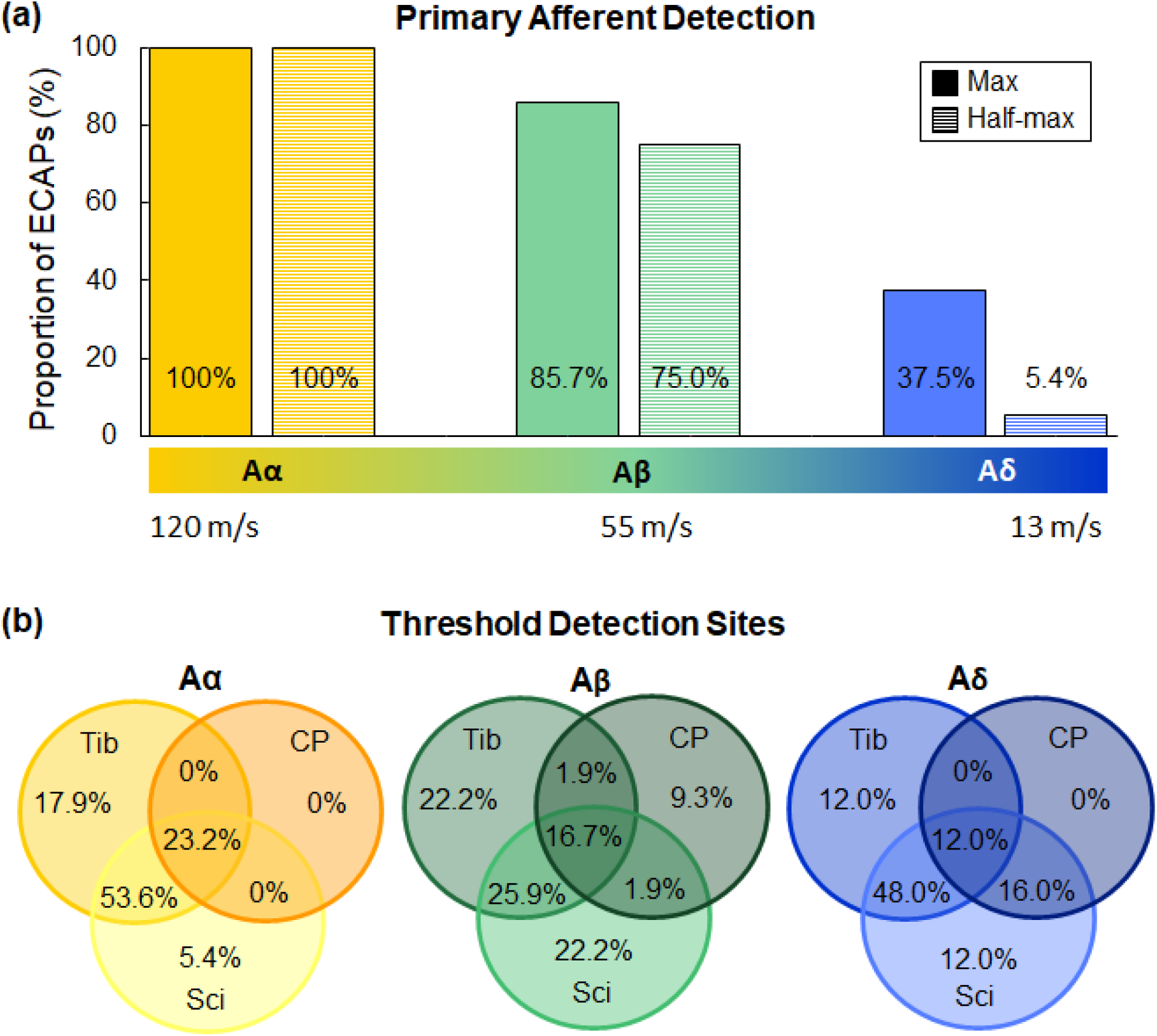
Detection of evoked compound action potentials (ECAPs) from primary afferents using both electrode types. (a) Proportion of ECAPs from each primary afferent (Aα, Aβ, and Aδ) at the maximum stimulation amplitude (just below the motor threshold; solid) and at half-max (the stimulation amplitude midway between threshold and the maximum; horizontal bars). (b) Proportion of Aα, Aβ, and Aδ thresholds that were detected at each nerve. Tib = tibial nerve; CP = common peroneal nerve; Sci = sciatic nerve. Where the circles intersect and overlap, the response at threshold were detected simultaneously on multiple cuff electrodes.

### ECAP thresholds for all afferents were most often recorded from the sciatic and tibial nerves

In order to compare the activation profiles of specific nerves with DRG stimulation using each electrode type, we also examined the nerve recording site from which we detected the ECAP response at threshold for each fiber type (Figure 6b). We detected Aα fiber ECAPs in only the tibial or sciatic nerves 17.9% and 5.4% of the time, respectively. At threshold, we detected a majority of Aα fiber ECAP responses in both the tibial and sciatic nerves (53.6%). At threshold, we did not detect Aα fiber ECAP responses from the common peroneal nerve, except when the response occurred simultaneously at all three nerve sites, which occurred in 23.2% of the thresholds detected. We detected ECAP thresholds for Aβ fibers in each of the three nerves. At threshold, we detected responses from Aβ fibers from only one of the tibial, common peroneal, or sciatic nerves 22.2%, 9.3%, and 22.2% of the time, respectively. Similar to the Aα fiber responses, we detected many Aβ fiber responses in both the tibial and sciatic nerves (25.9%), and seldomly from the common peroneal nerve (common peroneal and tibial = 1.9%; common peroneal and sciatic = 1.9%). At threshold, we detected some Aβ fiber responses from all three nerves (16.7%). The ECAP thresholds for Aδ fibers followed a similar trend, in which almost half of the responses that we detected were from both the tibial and sciatic nerves (48.0%), and approximately a quarter from each of those nerves separately (tibial = 12.0%; sciatic = 12.0%). We did not record responses from Aδ fibers from the common peroneal nerve alone, but 16.0% of the responses that we detected were from both the common peroneal and sciatic nerves, and 12.0% of the responses from all three nerves. In summary, we recorded ECAP responses at threshold most often from the sciatic and tibial nerves, either individually or in combination, for all three primary afferent fiber types.

### Injectrodes and stainless steel electrodes have similar recruitment rates that decrease as pulse width increases

We constructed recruitment curves for Aα and Aβ fibers by plotting the peak-to-peak amplitude of the ECAP versus the stimulation charge (current amplitude × pulse width) for each pulse width for the Injectrode and stainless steel electrode. Representative recruitment curves for Aα and Aβ fibers recorded from cat D with stimulation on the L6 DRG are shown in Figure 7a. The separation of the curves by threshold between the stainless steel electrode and Injectrode is consistent with the trends and differences noted in the group thresholds for L6 (Figure 4a-b).

**Figure 7.**
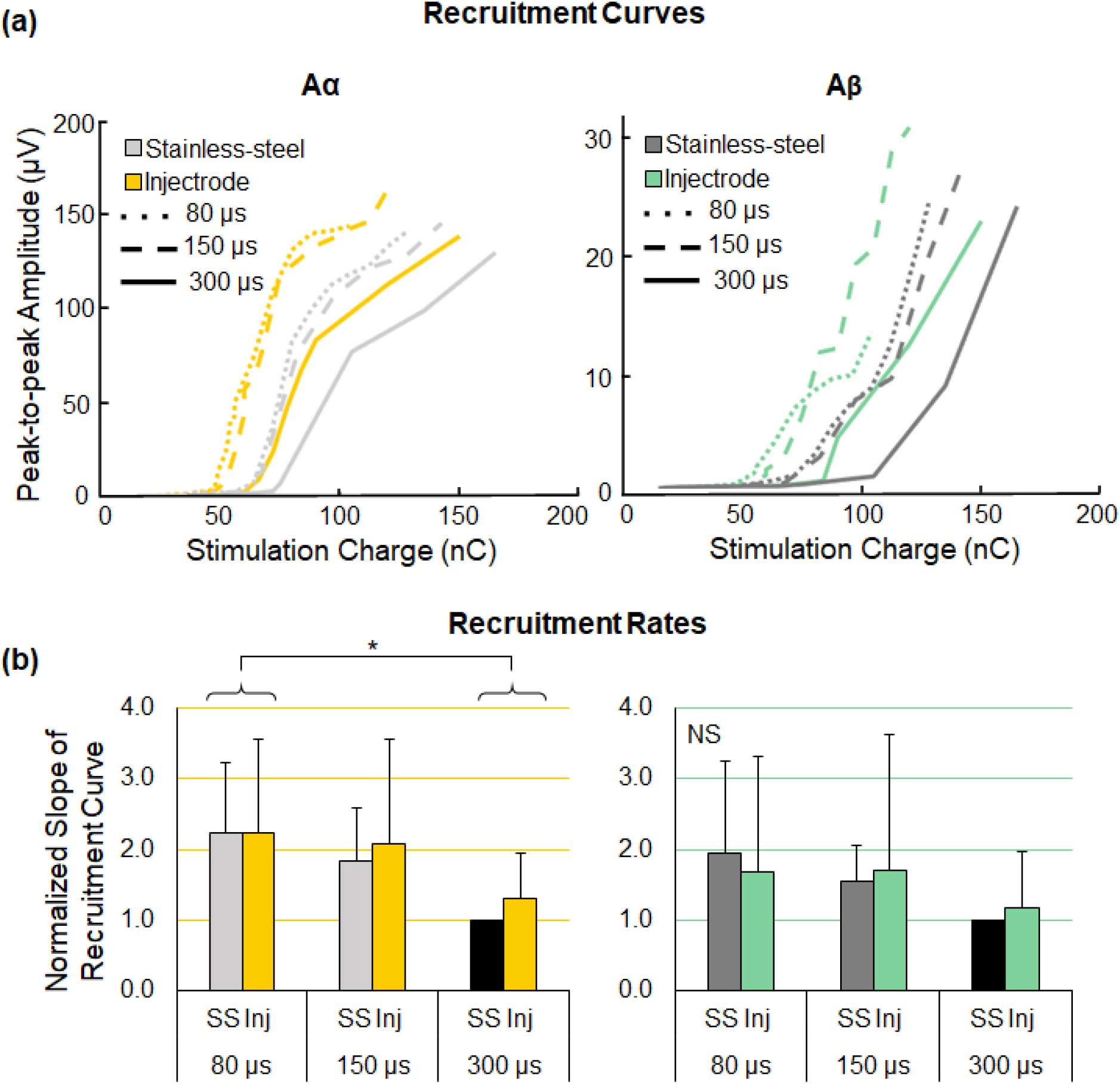
Recruitment properties of the evoked compound action potentials (ECAPs). (a) Recruitment curves for Aα and Aβ fibers showing the peak-to-peak amplitude of the ECAPs as a function of stimulation charge, for the stainless steel (SS) electrode and Injectrode (Inj) and across stimulation pulse widths. Representative data are from cat D, L6. (b) The rate of recruitment is the slope of the linear region of the recruitment curve. For each condition, the slope was normalized to the slope from the stainless steel electrode at 300 μs (black). *p ≤ 0.05; NS = not significant.

We defined the recruitment rates for Aα and Aβ fibers as the slope of the linear region of the recruitment curves. The slopes were normalized to the slope of the recruitment curve for the stainless steel electrode and a pulse width of 300 μs. For both Aα and Aβ fibers, the slope of the recruitment curves generally decreased as pulse width increased (Figure 7b); however, this difference was only significant for the Aα recruitment curves at 80 μs versus 300 μs (p = 0.011). The recruitment rates for the Injectrode were not statistically different from the recruitment rates for the stainless steel electrode across pulse widths for both Aα fibers (slope_INJECTRODE_ = 1.69 ± 0.87; slope_STAINLESS_STEEL_ = 1.87 ± 1.22; p = 0.54) and Aβ fibers (slope_INJECTRODE_ = 1.50 ± 0.87; slope_STAINLESS_STEEL_ = 1.52 ± 1.47; p = 0.95). Additionally, there were no interaction effects between the electrode type and pulse width (Aα: p = 0.89; Aβ: p = 0.87).

### Injectrodes have similar impedances that are lower than stainless steel electrodes

We measured EIS of the Injectrode and stainless steel electrode in cats C and D before stimulation began. The stainless steel electrode used in both cats was the same; however, the Injectrodes were freshly mixed and injected for each experiment. The stainless steel electrode and the Injectrodes had similar 1 kHz impedance magnitudes (stainless steel |Z| = 0.19 kΩ, 0.19 kΩ; Injectrode |Z| = 0.30 kΩ, 0.36 kΩ). Higher-frequency impedances (> 1 kHz) were similar between the two electrode types, indicating a similar electrolyte resistance (Figure 8a-b). At lower frequencies (< 100 Hz), the Injectrode had a smaller impedance magnitude and became less capacitive (impedance phase approached zero) at a lower frequency value than the stainless steel electrode. In cat D, we also collected EIS data after stimulation trials were completed for three separate Injectrodes placed on different DRG (Figure 8c-d). Following thousands of stimulation pulses over several hours (between 6 and 11 hours), the three different Injectrodes had similar EIS profiles across the frequency spectra. Notably, the impedance magnitude of the Injectrodes at 1 kHz remained below 1 kΩ after stimulation (|Z|_1_ = 0.26, |Z|_2_ = 0.41, |Z|_3_ = 0.35 kΩ). These values were also similar to the 1 kHz impedance of the Injectrodes prior to stimulation.

**Figure 8.**
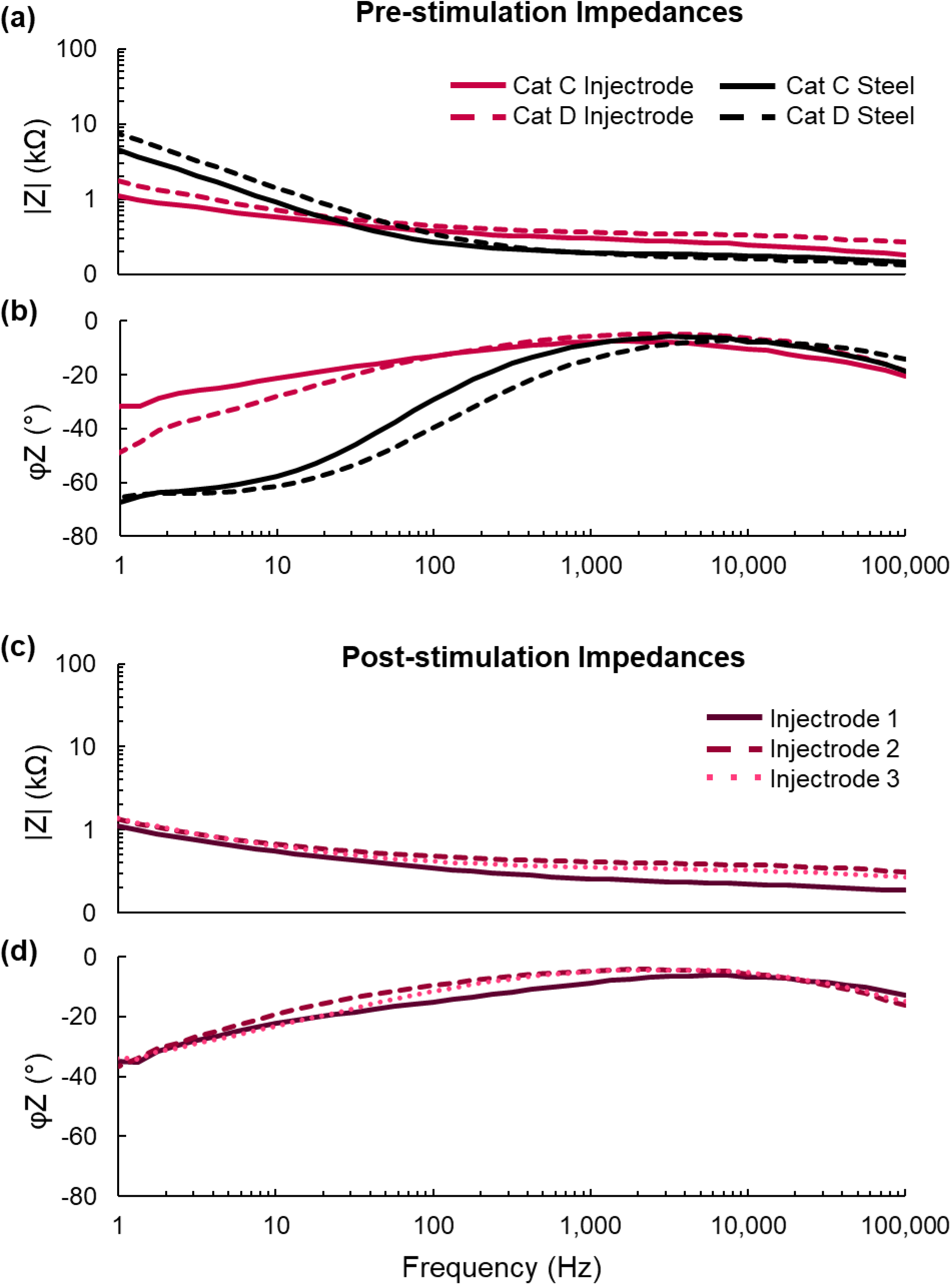
Electrochemical impedance spectroscopy. Impedance magnitude (a) and phase (b) for the stainless steel electrode and Injectrodes (from cats C and D) before stimulation. Impedance magnitude (c) and phase (d) for three different Injectrodes (cat D) after stimulation.

## DISCUSSION

### The Injectrode^®^ for DRG Stimulation

We compared the ECAP threshold and recruitment rate between the Injectrode and stainless steel electrode across several pulse widths. Data for stimulation of the L6 versus L7 DRG were analyzed separately because there were significant differences between the ECAP thresholds for Aα and Aβ fibers between the levels. Differences for Aδ fibers were not detected between levels but may be due to the small number of ECAPs from these small diameter fibers. Different levels of the DRG have different distributions of cell bodies and axons (Sperry et al., 2020), which may affect recruitment.

Our results show that the threshold for eliciting ECAPs using the Injectrode was not different or was significantly lower than the ECAP threshold using the stainless steel electrode (Figure 4). The average surface area of the Injectrodes were 1.96 times larger than the surface area of the stainless steel electrode. If the Injectrode and stainless steel electrode had a similar threshold, which was the case most of the time, then the Injectrode had a charge density at threshold that was 1.96 times lower than the stainless steel electrode. When the Injectrode had a lower threshold than the stainless steel electrode, which occurred for Aβ fibers at L6, then the Injectrode had an even smaller charge density compared to the stainless steel electrode. These approximations are based on the geometric surface area of the Injectrode; however, because the Injectrode is a polymer, it has a high porosity (Trevathan et al., 2019). A high porosity increases the permeability of the electrolyte, which increases the effective electrochemical surface area and likely reduces the charge density further. The similar-to-lower charge density of the Injectrode, as compared to the stainless steel electrode, is beneficial in several ways. First, it highlights the benefits of using a soft, flexible electrode that conforms to the DRG. Due to the small sub-laminar space in the cat model, the Injectrode made close-contact with the DRG, which increased the contact area between the Injectrode and the neural tissue. When an electrode is placed closer to neural tissue, it is able to excite more neurons at the same charge density (Gaines et al., 2018). Because the thresholds using the Injectrode were not different or smaller than the thresholds for the stainless steel electrode, resulting in a similar or lower charge density, stimulation through the Injectrode may reduce irreversible electrochemical reactions and the generation of toxic by-products that are harmful to tissue (Cogan, 2008). The Injectrodes also showed consistency over time: the ECAP thresholds measured at two timepoints separated over several hours were equivalent within a ±40% window. Consistent ECAP thresholds suggest that the cured Injectrode was able to maintain its material properties, such as high porosity, after several hours of interfacing with the tissue, as well as after thousands of stimulation pulses.

We measured recruitment of Aα, Aβ, and Aδ fibers with stimulation amplitudes below motor threshold, and Aα and Aβ fibers were recruited at thresholds much lower than Aδ fibers. We detected responses from large-diameter fibers in the majority of trials at maximum stimulation amplitude (Aα, 100%; Aβ, 85.7%) and at half-maximum amplitude (Aα, 100%; Aβ, 75.0%); however, responses from small-diameter Aδ fibers were identified in only 37.5% and 5.4% of trials at maximum and half-maximum, respectively. Our data, in agreement with previous studies (Bourbeau et al., 2011; Fisher et al., 2014; Gaunt et al., 2009; Nanivadekar et al., 2019), showed that DRG stimulation preferentially recruits large and medium diameter afferents, including Aβ fibers purported to promote analgesia according to the gate control theory of pain (Chao et al., 2020; Graham et al., 2019; Kent et al., 2018). Here, we show that the Injectrode is equally or more effective at recruiting Aβ fibers as the stainless steel electrode, highlighting that the Injectrode would be an appropriate electrode choice for DRG stimulation and treating chronic pain.

Avoiding activation of small-diameter fibers is ideal because some of these fibers transmit nociceptive signals that can cause pain. We stimulated the DRG across a range of amplitudes from subthreshold to relatively high amplitudes (up to 3.4 mA; motor threshold) in order to generate the recruitment curves. In preclinical studies, motor threshold is generally used as the upper bound of the amplitude range for stimulation (Chao et al., 2020). Clinical DRG stimulation is generally delivered at a current level that evokes paresthesias, indicative of Aβ fiber activation, but does not generate muscle activity (Kramer et al., 2015; Liem et al., 2013; Mekhail et al., 2020; Van Buyten et al., 2015). Recent modeling studies suggest that amplitudes greater than 4 mA are required to activate C fibers in human DRG (Graham et al., 2019; Kent et al., 2018). This amplitude is greater than the thresholds for evoking responses in Aβ fibers with a 300 μs pulse width using the Injectrode (331.7 μA) and stainless steel electrode (428.6 μA), as well as the typical clinical range of 1 mA or lower (Deer et al., 2017). We detected responses from Aδ fibers infrequently and at much higher thresholds than Aα and Aβ fibers, which is supported by computational models (Graham et al., 2020). Therefore, using the Injectrode for DRG stimulation with parameters that are used clinically should result in minimal activation of small-diameter fibers.

We measured recruitment thresholds using stimulation pulse widths of 80, 150, and 300 μs. Our data also showed that ECAP thresholds increased as the pulse width became longer in duration (Figure 4a-c). Although our trends were not always significant, likely due to being underpowered, this result is expected based on known charge-duration curves (Durand, 2006; Merrill et al., 2005). Clinical studies of DRG stimulation have reported mean pulse widths of approximately 300 μs (Deer et al., 2017; Graham et al., 2019; Kent et al., 2018; Liem et al., 2013) with the median pulse width decreasing to 255 μs at one year post-implant to reduce paresthesias (Deer et al., 2019). Our data suggest that it may be worthwhile to explore even shorter pulse widths clinically, as they may achieve comparable levels of Aβ fiber recruitment at lower charge levels. This is consistent with findings reported from a computational modeling study of DRG stimulation (Graham et al., 2020). Reducing charge delivery would also extend the battery life of DRG implants.

Although only a small number of EIS measurements were performed in this study, the results indicate that the impedances of Injectrodes were lower than the stainless steel electrode (Figure 8a-b), which is likely due to the larger surface area and porosity of the Injectrode. The 1 kHz impedance of the Injectrodes was below 1 kΩ, which is consistent with the benchtop data (Trevathan et al., 2019). The high-frequency impedances were similar for the stainless steel electrode and the Injectrodes before stimulation. This result is to be expected since the high frequency spectra indicate the component of the impedance due to the electrolyte resistance (Cogan, 2008; Dalrymple, 2021), which in this case is the same tissue and saline environment. The low-frequency impedances are dominated by the properties of the electrode-tissue interface and demonstrate the standard constant phase element behaviour of solid metal electrodes (Cogan, 2008; Dalrymple et al., 2019; Lempka et al., 2009; Lisdat and Schäfer, 2008). The Injectrodes have a lower impedance that is more resistive and less capacitive than the stainless steel electrode, indicated by the lower impedance magnitude and an impedance phase that is closer to zero degrees. This is advantageous because it shows that the Injectrodes have less electrode polarization, or charge accumulation (Dalrymple et al., 2019). Pt electrodes, which are more clinically-relevant, also exhibit more capacitive behaviour at low frequencies, indicated by a phase closer to −90°. This behaviour has been reported in several *in vivo* studies using Pt electrodes, in which the low frequency (1 Hz) impedance phase was approximately −66° (Shepherd et al., 2021) and the 100 Hz impedance phase ranged from −56° to −72° (Dalrymple et al., 2020a, 2020b; Shepherd et al., 2021). This is similar to how the stainless steel electrode behaved here, suggesting that the Injectrode may also accumulate less charge than a Pt electrode. However, a direct comparison must and will be done in the future. The small low-frequency impedances of the Injectrodes are likely due to the porosity of the silicone matrix, making the silver particles more accessible to the ions in the electrolyte/tissue (Ludwig et al., 2006; Trevathan et al., 2019). The more resistive impedance is also characteristic of silver electrodes, which form silver chloride during stimulation (Trevathan et al., 2019).

The EIS traces for the three different Injectrodes following stimulation were similar across the frequency spectra (Figure 8c-d). This result is especially interesting for two reasons. First, each of these Injectrodes were delivered through a burr hole in the lamina. Therefore, we had no control over the shape and distribution of the Injectrode material within the intraforaminal space; only the volume of Injectrode was held constant. Secondly, EIS was recorded after stimulation was completed. The Injectrodes were implanted between 6 and 11 hours prior to the EIS recordings, and they were stimulated intermittently within that time. This result indicates that, at least in the acute setting, repeated stimulation through the Injectrode, and interaction with the tissue over several hours, does not induce a measurable change in the electrical properties of the Injectrode or its ability to stimulate nerve fibers.

### Limitations, Relevance, and Translation to Humans

ECAPs were recorded from the peripheral nerves in response to DRG stimulation in anaesthetized cats. Cats were chosen as a model because they are a common model for DRG stimulation, particularly by our group (Ayers et al., 2016; Fisher et al., 2014; Gaunt et al., 2009; Nanivadekar et al., 2019). Clinical stimulation typically uses a bipolar configuration, where two contacts are placed to straddle the DRG (Deer et al., 2019). We utilized monopolar stimulation to first ensure that DRG stimulation was possible with the Injectrode, and future iterations will include more flexibility with contact configurations. Nonetheless, our results showing that Aα and Aβ fibers exhibit the lowest thresholds for activation, which is consistent with previous findings using computational models (Bourbeau et al., 2011; Chao et al., 2020; Graham et al., 2019, 2020; Kent et al., 2018).

At ECAP threshold, the majority of responses were identified in the sciatic and tibial nerves. Despite the tibial and common peroneal nerves being distal branches of the larger sciatic nerve, ECAPs were not always detected in the sciatic nerve when detected in the smaller branches. This result has been reported previously and it likely occurs because the sciatic nerve is larger, making it more difficult to detect responses in smaller nerve fibers using the nerve cuff electrode (Ayers et al., 2016). We detected more than three-quarters of Aα ECAPs at threshold in either the tibial or sciatic nerve, while the remaining ECAPs were detected across all nerves (Fig. 6b). Responses in Aβ fibers were spread across all nerves, but less than 10% of ECAPs were identified only in the common peroneal at threshold (Fig. 6b). Similar to our findings, at threshold, Aα ECAP responses from a prior study using epineural stimulation of the L6 and L7 DRG were most often present in the sciatic and tibial nerves, both individually and in combination (Nanivadekar et al., 2019). Anatomical differences between the tibial and common peroneal nerves likely explain their varying patterns of activation by DRG stimulation. The tibial nerve is much larger than the common peroneal nerve, containing many more axons and thus increasing the likelihood of activation during stimulation (Apaydin and Bozkurt, 2015). The tibial nerve innervates the muscles and skin of the calf, lower leg, and foot, and plays a critical role in ankle control and stability during standing and walking (Akazawa et al., 1982; Capaday and Stein, 1986; Konradsen et al., 1993). Therefore, proprioceptive information (relayed by Aα fibers) from the ankle is important and may explain the prevalence and ease of detection of Aα ECAPs from the tibial nerve. The tibial and common peroneal nerves innervate the heel and dorsum of the foot, respectively. Mechanosensation from the foot, innervated by the tibial and common peroneal nerves, plays a large role in modulating gait and reflexive responses to perturbations (Van Wezel et al., 1997; Zehr et al., 1997). This anatomy may explain why we were able to detect ECAPs from Aβ fibers at threshold in each of the nerves (Bear et al., 2007).

The mechanisms of pain differ between males and females, as do their responses to therapies (Bartley and Fillingim, 2013; Catuneanu et al., 2019; Fillingim et al., 2009; Mogil, 2012). In the current study, all experiments were performed in male cats. We aimed to compare the recruitment properties of DRG stimulation using the Injectrode or a stainless steel electrode, which did not require us to evaluate a model of neuropathic pain. In the future, as we seek to validate the Injectrode in models of neuropathic pain, we will use both male and female animals.

We demonstrated that the Injectrode can be delivered onto the DRG using a needle and syringe. We were able to successfully target the DRG by injecting the Injectrode into a burr hole made in the lamina. We were able to identify the location of the DRG by using the exposed contralateral DRG as a guide, assuming symmetry about the spinal cord. However, fluoroscopic guidance could also be used to locate the target DRG, as is the standard practice for implanting electrodes for SCS and DRG stimulation. The burr hole exposure worked well for targeting the lumbar DRG in the cats because their DRG are close to the spinal canal and lie almost parallel to the spinal cord. Depending on the spinal level, the DRG in humans can lie either inside the spinal canal (sacral) or in the intraforaminal space (lumbar) (Haberberger et al., 2019; Kikuchi et al., 1994). We are currently exploring alternative Injectrode delivery approaches for translation to humans, including a transforaminal approach similar to what is used for steroid injections (Rivera, 2018). This type of approach has also been proposed as a method to locate the optimal DRG level using radio frequency stimulation (Hunter et al., 2017; Zuidema et al., 2014) and is used to target the sacral DRG (Deer et al., 2019). Additionally, one study has reported the use of a DRG stimulation system (StimWave Inc., Pompano Beach, FL, USA) implanted transforaminally in 11 subjects at the L1 to L5 DRG levels (Weiner et al., 2016). A transforaminal approach may be simpler than the current surgical approach for implanting DRG stimulation leads, as it would not require steering of the leads outside of the cannula into the foramen (Caylor et al., 2019; Deer et al., 2013; Liem, 2015).

The formulation of the Injectrode employed in this study had silver particles mixed within a polymer. In large quantities, silver and silver compounds are neurotoxic (Drake and Hazelwood, 2005; Lansdown, 2007). However, silver is a cost-effective conductive material, and was used in this proof-of-concept study to demonstrate that (i) the Injectrode can easily be delivered onto the DRG using a burr hole method; (ii) the Injectrode can effectively excite primary afferent neurons in the DRG; and (iii) the Injectrode has similar recruitment properties as a cylindrical stainless steel electrode. While clinical DRG leads are designed for chronic stimulation and made of platinum, which offers superior corrosion resistance (Cogan, 2008; Harnack et al., 2004; Stevenson et al., 2010; Wellman et al., 2018) and higher charge storage capacity (Merrill et al., 2005), the custom stainless steel electrode used in this study exhibited low impedance and stable performance over the relatively short duration of this study. Stainless steel electrodes are frequently used in electrical stimulation studies (Babb and Kupfer, 1984; Desai et al., 2014; Gordon et al., 2010; Popovic et al., 1991; Walter et al., 1993) because they are an effective alternative to more expensive metals used in clinical leads that are implanted chronically (ex. platinum).

Silver and other metallic particles, including gold and carbon black, are commonly used in the development of soft and flexible electronics (Araki et al., 2011; Cong and Pan, 2009; Larmagnac et al., 2014; Minev et al., 2015; Niu et al., 2007). In preparation for chronic studies, and the eventual translation to human implants, we are exploring other Injectrode formulations that use gold or platinum particulates, as well as other Injectable electrode designs that are wire-based, which would be easier to remove if needed. Gold or platinum are more appropriate than silver for chronic implants because they are biocompatible and are less likely to elicit a strong tissue response (Wellman et al., 2018). A strong tissue response would increase the impedance at the electrode-tissue interface, thereby requiring more charge injection to effectively stimulate the neural target (Cogan, 2008; Dalrymple, 2021; Dalrymple et al., 2020a). The Injectrode design is promising for chronic use as it conforms to the tissue around it, limiting the potential for compression of nearby structures, like the DRG, as compared to stiffer electrode leads currently used for DRG stimulation. In-body curing polymers are being explored for other uses as well, including for aneurysm grafts but have been limited to *in vitro* and human *ex vivo* studies (Bosman et al., 2010; van der Steenhoven et al., 2012). Other types of injectable electrodes are also under investigation in animal models, such as an injectable mesh electronic probe used to record neural activity from the rat basal ganglia (Schuhmann et al., 2017). Collectively, the recent developments towards soft, flexible, in-body curing electrodes aim to reduce the invasiveness and stiffness of implanted electrodes and, ultimately, reduce the biotic reactions and extend the lifetime of neural interfaces. So far, the Injectrode has been used to stimulate the brachial plexus in rats and the swine vagus nerve (Trevathan et al., 2019), and now, the feline DRG. Our studies demonstrate that the Injectrode has a wide range of potential uses in the field of neurostimulation. An improvement to the Injectrode design that is currently underway includes wireless stimulation, which will reduce the invasiveness and allow more flexibility with programming and power supply upgrades.

Further success of the Injectrode for neurostimulation applications can be achieved by creating a multi-contact Injectrode. Typically, DRG stimulation utilizes bipolar stimulation, where the best results occur when two electrode contacts straddle the DRG (Deer et al., 2019; Eldabe et al., 2015; Graham et al., 2019) due to maximal Aβ fiber activation (Graham et al., 2020). Furthermore, multi-contact electrode arrays allow for more options for electrode selection and target areas over the neural target. This capability is especially useful for expanding the use of the Injectrode to other neuromodulatory applications, such as SCS, and will be explored in future work.

## CONCLUSIONS

We stimulated the feline L6 and L7 DRG with either the Injectrode or a cylindrical stainless steel electrode to evaluate the Injectrode as an effective device for DRG stimulation. We exposed the DRG via a partial laminectomy or a burr hole. Overall, DRG stimulation using the Injectrode had a similar or lower threshold to evoke compound action potentials in Aα, Aβ, Aδ fibers than a cylindrical stainless steel electrode; thresholds increased as pulse width increased. Thresholds for the Injectrode were consistent over several hours and thousands of stimuli. The rates of recruitment for Aα and Aβ fibers were not different between the Injectrode and stainless steel electrodes and generally decreased as pulse width increased. Injectrodes had lower or similar impedances relative to the stainless steel electrode across frequency spectra that were consistent after stimulation, despite having cured with slightly different conformations following injection through the burr hole. The Injectrode is less-stiff than traditional DRG electrodes and can conform to the shape of the implant target and surrounding cavity. These properties may improve the recruitment of Aβ fibers and enhance the stability of DRG stimulation to provide pain relief to patients suffering from chronic pain.

## ACKNOWLEDGEMENTS

We would like to thank the Staff at Magee Women’s Research Institute Animal Facility and Rachel Pitzer for their assistance with animal care and monitoring. We would also like to thank Ritesh Kumar for his technical assistance, as well as the Cohen-Karni lab at CMU for lending us the electrodes for performing the electrochemical measurements.

